# Mendelian Randomization analyses reveal a causal effect of thyroid function on stroke via atrial fibrillation

**DOI:** 10.1101/718429

**Authors:** Eirini Marouli, Aleksander Kus, M. Fabiola Del Greco, Layal Chaker, Robin Peeters, Alexander Teumer, Panos Deloukas, Marco Medici

## Abstract

**Background:** Several observational studies suggest that variations in thyroid function, even within the normal range, are a risk factor for cardiovascular diseases, but it remains to be determined if these associations are causal or not. This study investigates whether the relationship between variation in normal range thyroid function, as well as hypothyroidism and hyperthyroidism, and the risk of stroke and Coronary Artery Disease (CAD) are causal and via which pathways these relations are mediated.

**Methods and Findings:** We performed Mendelian Randomization (MR) analyses for stroke and CAD using genetic instruments associated with TSH and FT4 levels respectively within either the normal range, hypothyroidism or hyperthyroidism. In detected associations, the potential mediatory role of known stroke and CAD risk factors was also examined. A one standard deviation increase in TSH was associated with a 5% decrease in the risk of stroke (OR=0.95, 95% CI= 0.91 to 0.99). Multivariable MR analyses indicated that this effect is mediated through atrial fibrillation (AF). Hashimoto’s Disease (HD) was associated with a 7% increased risk of CAD (OR=1.07, 95% CI= 1.01 to 1.13). The effect of Hashimoto’s Disease (HD) on CAD risk appears to be mediated via body mass index (BMI).

**Conclusions:** These results provide important new insights into the causal relationships and mediating pathways between thyroid function, stroke and CAD. Specifically, we identify normal range TSH levels and HD as potential modifiable risk factors for stroke and CAD, respectively.

## Introduction

Despite the undeniable progress in prevention and treatment in the past two decades, cardiovascular disorders (CVD) remain the leading cause of mortality worldwide (1). Whereas smoking, hypertension, diabetes, obesity and dyslipidaemia are the major modifiable cardiovascular risk factors(2), observational studies have demonstrated that also overt and subclinical thyroid dysfunction are associated with a higher risk of CVD(3–6). Moreover, even variation in thyroid function within the normal range has been associated with an increased risk of CVD, including atherosclerotic disease and stroke (7–10). This poses the question as to whether common forms of mild thyroid dysfunction (with a prevalence of 5-10 % in the general population) should be treated to prevent these complications. Unfortunately, no data on the effects of treatment of these milder forms of thyroid dysfunction on cardiovascular mortality or morbidity are available from well-powered randomized controlled trials (RCTs) (11, 12). Moreover, given the follow-up time and sample sizes required for such conclusive RCTs, these data will not become available in the next few years (if ever). Mendelian randomization (MR) is another approach that can provide crucial information on causality when RCTs are not feasible or unavailable (13). MR evaluates the effect of an exposure (e.g. thyroid function) on an outcome (e.g. CVD) using genetic variants associated with the exposure as instruments(14). MR draws from the fact that genetic variants segregate randomly from parents to offspring, which is comparable to the randomization used in clinical trials. As 65% of the total variance in TSH and FT4 levels is determined by genetic factors (15), there are good grounds for MR studies on thyroid function and various outcomes. Evidence from a previous MR study suggests no causal association between thyroid traits (TSH, FT4) and ischemic heart disease(16). However, this study had only limited power due to the small number of genetic variants (variance explained: 5.6% for TSH and 2.3% for FT4) used as instruments (17).

Recently, the ThyroidOmics consortium reported the largest meta-analysis of genome-wide association studies (GWAS) on thyroid function in over 72,000 subjects which more than doubled the number of loci associated with thyroid function (18). These findings paved the way to conduct well-powered MR studies to test the causality of the observed associations between thyroid function and CVD. To this end, we performed two-sample MR to investigate the role of thyroid function on CAD and stroke respectively using as instrument the variants reported from the above GWAS on thyroid function and publicly available summary statistics from the two largest GWAS on CAD(19) and stroke(19, 20). Next to normal range thyroid function, MR studies on subclinical and overt thyroid disease, including Hashimoto’s disease (HD) and Graves’ disease (GD), are presented as secondary analyses, thereby covering the entire spectrum of thyroid (dys) function. Finally, mediation analyses were performed to investigate the pathophysiological mechanisms underlying the causal associations.

## Methods

### Two-sample Mendelian Randomization

We performed two-sample MR analysis by using summary genetic data from the largest GWAS studies available for thyroid function (18), CAD(19) and stroke(20). The exposures of interest included: normal range TSH and FT4 levels, GD and HD. As subclinical hypo-and hyperthyroidism have been associated with CAD and stroke, we additionally investigated increased TSH and decreased TSH levels, respectively. Summary data of the variants identified to be associated with the exposure were extracted. CAD data were derived from the largest systematic GWAS meta-analysis, involving 122,733 CAD cases and 424,528 controls from the van Harst *el al* (19). Stroke data were derived from the recent multi-ethnic meta-analysis for any stroke, provided by the MEGASTROKE consortium (20).

The two-sample MR was performed using the Inverse Variance Weighted (IVW)(21) method (assuming that all the IV assumptions are met). We also used other methods relaxing the third assumption about pleiotropy (MR-Egger (Egger)(22); Weighted Median (WM)(23)). An IVW corrected for the SEs of each instrument, for the correlation between the associations of the instrument with the exposure and the association of the instrument with the outcome was also used. Heterogeneity for the Wald ratios was tested with the Cochran’s Q and quantified with the I^2^ index (24). The strength of the association between the exposure and the outcome was estimated by calculation of the F-statistic (25).

In order to quantify the strength of the “NO Measurement Error” assumption (NOME) for MR-Egger we used the I^2^ statistic of the genetic association estimates on the exposures. This measure is called I^2^_GX_ and lies between 0 and 1. A high value of I^2^_GX_ (close to 1) indicates that dilution bias does not affect the standard MR-Egger analyses performed (24). In the main analyses we used the first order weights, which correspond to the first term of the Taylor series expansion, which approximate the SE of the Wald ratio estimate (24). The IVW and the MR-Egger regression analyses were repeated using the second order weights, which correspond to the first two terms of the Taylor series expansion (24).

We also applied other MR methods relaxing the InSIDE (Instrument Strength Independent of Direct Effect) assumption (Mode-Based Estimate (MBE)(26)), relaxing the independence among IVs (Generalised *Summary*-data-based Mendelian Randomization (GSMR)(27)) and assuming that at least 50% of the IV are valid instruments (MR-PRESSO(28)) approaches, using summary level data (STable 14).

### Sensitivity Analyses

Sensitivity analyses were performed in order to further elucidate the role of pleiotropy in the association of thyroid function and disease. The Q statistic was used to assess variants that may possibly contribute to the total heterogeneity. Different thresholds were used (5th (L1), 1st (L2), 0.19th (L3) percentile of a chi-squared with 1 degree of freedom). Any variants which had a Q>L3, Q>L2 and Q>L1 were excluded(29).

### Genetic variants used as instruments

We used 55 and 29 single nucleotide polymorphisms (SNPs) associated at a genome-wide significant level (p<5e-8) with TSH and FT4 levels within the reference range, respectively, as instruments to investigate the causal role of minor variation in thyroid hormones levels on CAD and any stroke.

In order to cover the entire spectrum of thyroid (dys)function, for secondary analyses we assessed the causal effect of mild thyroid dysfunction on these two outcomes by using variants and summary data derived from two separate case–control GWAS meta-analyses of increased TSH levels (i.e., (subclinical and overt) hypothyroidism) and decreased TSH levels (i.e., (subclinical and overt) hyperthyroidism), including cases with TSH levels above and below the cohort-specific reference range, respectively, and controls with TSH levels within the reference range(18). One condition of MR is that exposure-related SNPs (the instrumental variables) must not be in Linkage Disequilibrium (LD) with each other, as that can result in confounding (30). For each trait, only the SNPs with the lowest association p-value that passed the PLINK LD pruning algorithm (31) was kept (1000Gv3 ALL samples was used as LD reference, r^2^≤0.01 within windows of ±1 Mb) (STables 17, 18).

We also performed a literature search for HD (20 variants) and GD (49 variants) associated variants to assess the causal role of thyroid disease on CAD and stroke. The summary statistics for these variants and their association with the corresponding thyroid disease were derived from UK Biobank.

### Analyses using UK Biobank data

The UK Biobank (UKBB) study recruited more than 500,000 individuals across Great Britain from 2006-2010(32)(http://biobank.ctsu.ox.ac.uk).

ICD10 codes used to define HD were E03.8, E03.9 and E06.3. For GD these were E05.0, E05.8, E05.9.

### Datasets

Summary statistics for TSH, FT4, increased and decreased TSH associated variants were extracted from the currently largest GWAS meta-analysis for thyroid function (18). For HD and GD risk variants we used summary associations derived from UKBB, as described above. CAD data were extracted from the most recent meta-analysis that used data from the Harst *el al.* meta-analysis(19), stroke(20), Type 2 Diabetes (T2D) from DIAGRAM(33), High Density Lipoprotein (HDL), Low Density Lipoprotein (LDL), total cholesterol (TC) and triglycerides (TG) were obtained from GLGC (http://csg.sph.umich.edu/abecasis/public/lipids2013/) and ENGAGE consortia (http://diagram-consortium.org/2015_ENGAGE_1KG/); anthropometric traits including Body Mass Index (BMI) from the GIANT consortium (https://portals.broadinstitute.org/collaboration/giant/index.php/GIANT_consortium_data_files); blood pressure (BP) traits from ICBP(34). HD and GD summary data were derived from UKBB(32). The age completed full time education were extracted from http://www.nealelab.is/uk-biobank. Atrial fibrillation (AF) data were extracted from Nielsen, J.B. *et al.* (35) and heart rate (HR) data from Verweij, N *et al.* (36).

Summary data including effect/other alleles, allele frequencies, beta coefficients, standard errors (SE) and p-values were extracted. Effect sizes were aligned to the increasing alleles for TSH or FT4 levels and to those that increase the risk of increased TSH levels, decreased TSH levels, hypothyroidism or hyperthyroidism. No ethical approval was required as all data were extracted from publically available summary data.

### Bayesian model averaging (MR-BMA) method

MR-BMA can detect true causal risk factors even when the candidate risk factors are highly correlated. It performs risk factor selection and determines which risk factors are the causal drivers on disease risk. This approach is then prioritising and ranking all potential mediating factors by their Marginal Inclusion Probability (MIP). We can further identify all models with a posterior probability > 0.01 and check their model fit. Cook’s distance or Q-statistic can point to variants as influential points. We repeat MR-BMA analysis without the variants presented as influential points and evaluate the top risk factors. We further evaluate any observation that has a consistently large Q-statistic or Cook’s distance(37).

### Multivariable Mendelian Randomisation Analyses

When MR showed a causal relationship between thyroid function and CAD or stroke, multivariable MR analyses were performed to evaluate the mediating role of possible risk factors. These mediators included: BP (Systolic BP (SBP), Diastolic BP (DBP), Mean Arterial Pressure (MAP), Pulse Pressure (PP)), T2D, BMI and lipid traits (TC, LDL, HDL, TG), as well as AF, heart rate (HR), educational attainment (EA))(assessed by age on which full time education was completed) using summary data. In these analyses, the proportion of the effect that may be mediated by another factor was evaluated by the change in the total effect of the genetically determined exposure on the outcomes(38). We additionally applied a Bayesian model averaging (MR-BMA) approach to multivariate MR in order to prioritise risk factors for CAD and stroke and to determine which out of a set of related risk factors with common genetic predictors are the causal drivers of the disease risk(37).

For information on datasets, participants, genetic variants, diagnoses of HD and GD and statistical analyses, see Supplementary Material online, Methods.

### Power calculations

To estimate the power of our study, we calculated minimal odds ratio (OR) of the outcome variable (CAD and stroke) per standard deviation (SD) of the exposure variable (TSH and FT4 levels) detectable (power = 0.8, α = 0.05) in our study, using a non-centrality parameter-based approach (39), implemented in a publicly available mRnd web tool (http://cnsgenomics.com/shiny/mRnd/). Proportion of total variance in TSH and FT4 levels explained by the genetic variants used as instruments (9.4% and 4.8%, respectively) was established based on the data from Teumer *et al* (18).

## Results

### Normal range TSH and FT4 levels

MR analyses indicated a significant causal association between higher TSH levels within the reference range and a lower risk of stroke (OR=0.95, 95% CI= 0.91- 0.99, p =0.008 per 1 SD increase in TSH levels). There was no evidence of directional pleiotropy (Egger intercept= 1.06e-5, SE= 0.003) (Fig 1A). Next to the IVW method, we applied additional MR approaches (see methods), as no single method controls for all statistical properties that may impact MR estimates. The GSMR and MR PRESSO analyses yielded similar results to IVM (STable14 A, B), further corroborating the causal association between higher TSH levels and a lower risk of stroke. The WM approach yielded marginally significant results, but thedirection and magnitude of the effect was the same as observed in the other methods (OR=0.95, 95% CI= 0.92 to 0.98, p =0.092) (STable 1).

**Fig 1.**
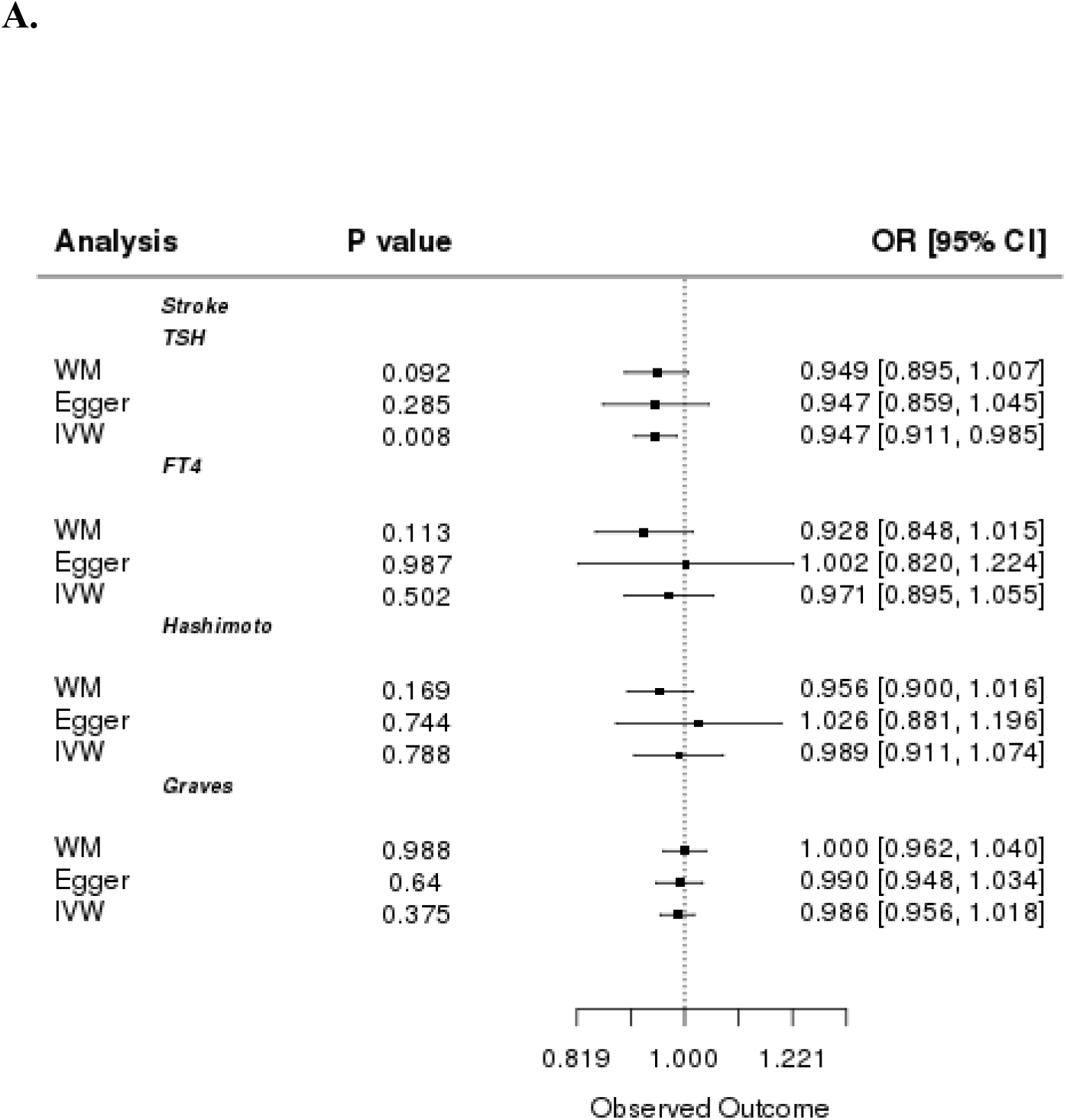
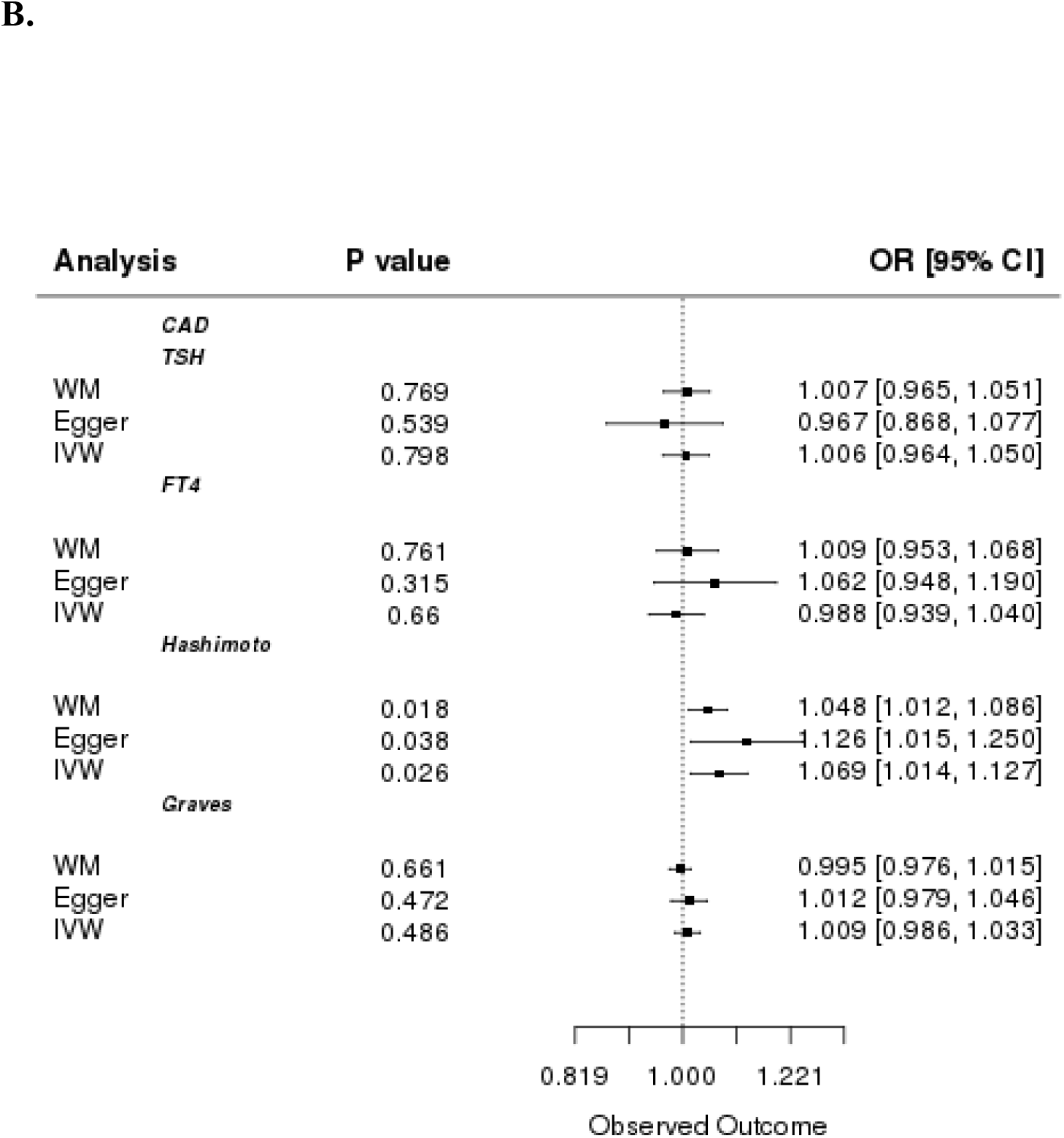
Two sample Mendelian Randomisation analyses - Estimates of the Effect of TSH, FT4 levels, Hashimoto’s and Graves’ Disease on A. Stroke and B. Coronary Artery Disease. IVW-Inverse Variance Weighted, MR-Egger, WM: Weighted Median Effect estimates represent the ORs (95% CI)

There was no evidence for a causal association between normal range FT4 levels and the risk of stroke (IVW: OR=0.97, 95% CI= 0.89- 1.06, p =0.502 per 1 SD increase in FT4 levels) (STable 2, Fig 1A)).

For CAD, no causal associations were detected with normal range TSH (OR=1.01, 95% CI= 0.96 to 1.05, p =0.8 per 1SD increase in TSH levels) or FT4 levels (OR=0.99, 95% CI= 0.94 to 1.04, p =0.66 per 1SD in FT4 levels) using IVW (STables 3 - 4). Similar results were obtained with the other methods (Stables 3-4, Fig 1B). Egger regression did not provide any evidence for directional pleiotropy (TSH: Intercept= 0.003, 95% CI= −0.004 to 0.010, FT4: Intercept= −0.005, 95% CI= −0.014 to 0.003).

### Hypothyroidism (increased TSH and Hashimoto’s Disease)

A significant positive causal effect of HD on the risk of CAD was observed using IVW (OR=1.07, 95% CI= 1.01 to 1.13, p =0.026) and corroborated by other methods (STable 5). Egger regression did not provide evidence for directional pleiotropy (Intercept= −0.008, 95% CI= −0.023 to 0.006) (STable 5).

There was no evidence for a causal effect of increased TSH on CAD risk (OR=1.04, 95% CI= 0.90 to 1.19, p =0.85, IVW) (STable 9). Similarly, no causal associations between increased TSH, HD, and stroke were detected (STables 6, 9).

### Hyperthyroidism (decreased TSH and Graves’ Disease)

No causal associations between GD and CAD (OR=1.01, 95% CI= 0.98 to 1.03, p =0.49, IVW) (STable 7) or stroke (OR=0.99, 95% CI= 0.96 to 1.02, p =0.37, IVW) were detected (STable 8). There was no evidence for causal associations between decreased TSH and stroke or CAD (STable 9).

## Mediation Analyses

To further investigate the observed causal association between normal range TSH levels and stroke, we assessed the role of potential stroke risk factors as mediators using multivariable MR analyses. Adjustment for the effect of each possible mediator showed that the effect of TSH levels on the risk of stroke disappeared after adjustment for AF (OR=1.00, 95% CI= 0.95 to 1.06, p=0.86). The observed causal effect in the unadjusted analyses was independent of BMI, T2D, lipids, BP, HR and educational attainment which was used as a surrogate for socio-economic status (STable 10A, Fig 2). To further assess the effect of TSH on the risk of stroke via AF, we investigated whether the known effect of TSH on AF (40) is mediated by any risk factors. Multivariable MR analyses did not implicate BP, lipids, CAD, or T2D as possible mediators of the observed causal effect of TSH levels on AF risk (STable 11).

**Fig 2.**
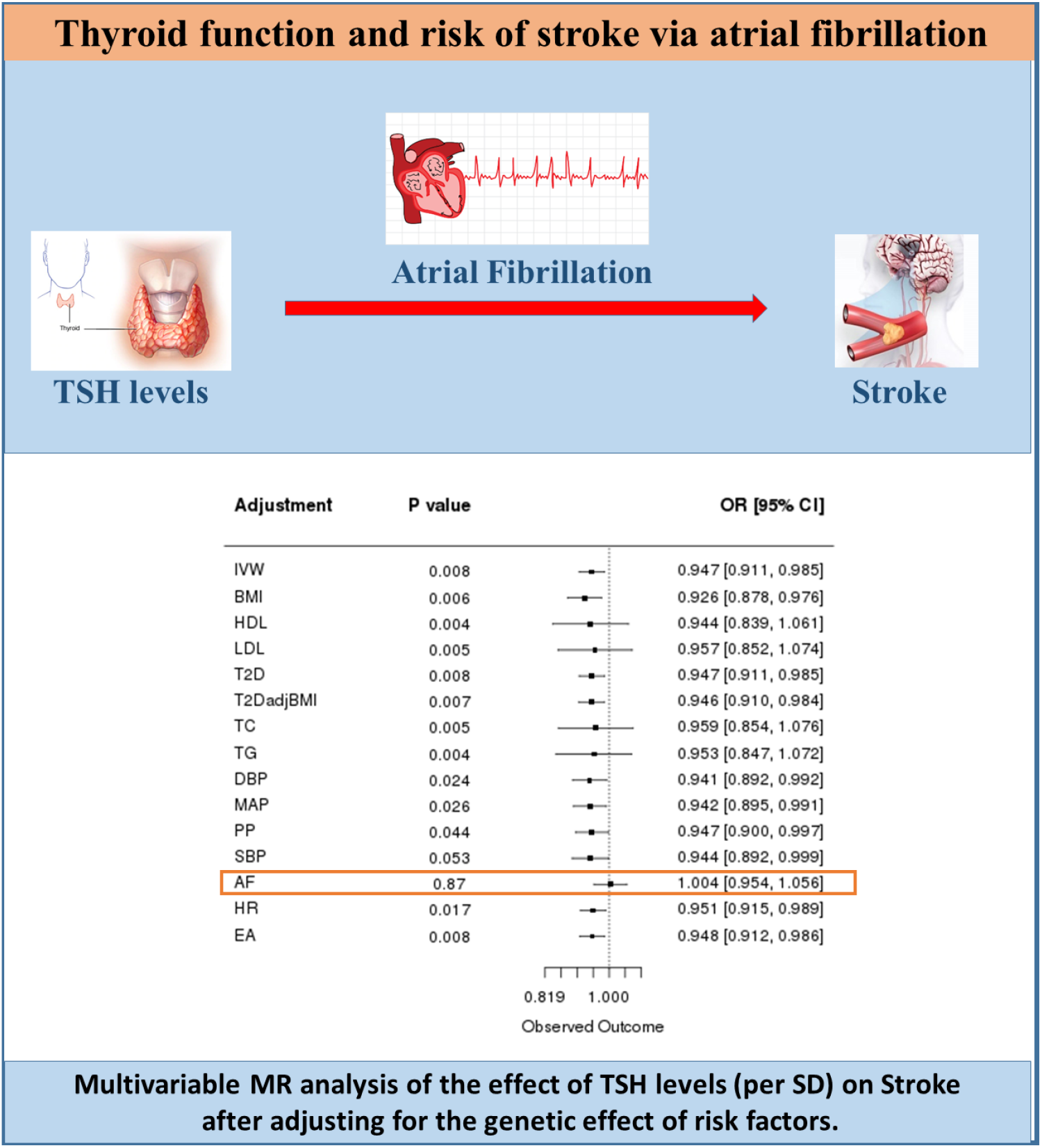
Multivariable MR analysis of the effect of TSH levels (per SD) on Stroke, after adjusting for the genetic effect of possible mediators. (MR-IVW: Mendelian Randomisation Inverse Variance Weighted, BMI: Body Mass Index, HDL: High Density Lipoprotein, LDL: Low Density Lipoprotein, T2D: Type 2 Diabetes, T2DadjBMI: Type 2 Diabetes adjusted for BMI, TC: total Cholesterol, TG: Triglycerides, DBP: Diastolic Blood Pressure, MAP: Mean Arterial Pressure, PP: Pulse Pressure, SBP: Systolic Blood Pressure, AF: Atrial Fibrillation, HR: Heart rate, EA: Educational attainment)

As the results presented above suggested that AF was a putative mediator for the effect of TSH levels on stroke risk, we used the MR-BMA method to further corroborate this finding. MR-BMA can detect true causal risk factors even when the candidate risk factors are highly correlated. This analysis also showed that AF was the top mediating factor (STable 12; SFig 1). Further inspection of the models indicated two variants (rs74804879, rs17477923) as influential points. MR-BMA analysis after excluding these two variants showed again AF as the top risk factor (STable 13).

Multivariable MR analyses were also performed for the observed causal association between HD and CAD to detect potential mediators. These analyses indicate that the effect of HD on CAD may be mediated via BMI (STable 10B; Fig 3). The MR-BMA method results are shown in STable 15, and one variant (rs10774625) was identified as influential point (STable 15C). MR-BMA analysis without this variant suggest BMI and HR as the most important risk factors in the relation between HD and CAD (STable 16).

**Fig 3.**
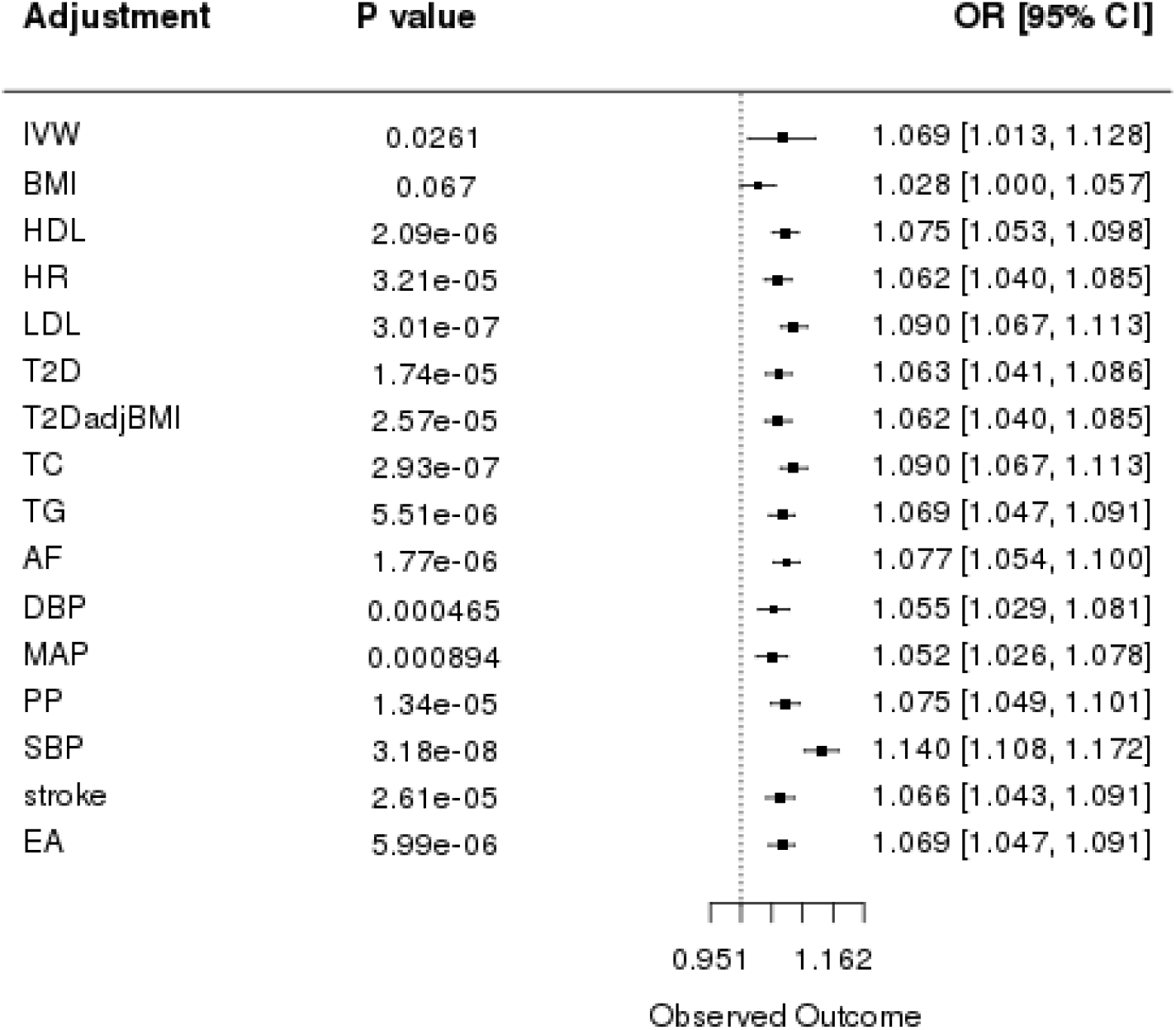
Multivariable MR analysis of the effect of Hashimoto’s Disease on CAD, after adjusting for the genetic effect of possible mediators. (MR-IVW: Mendelian Randomisation Inverse Variance Weighted, BMI: Body Mass Index, HDL: High Density Lipoprotein, LDL: Low Density Lipoprotein, T2D: Type 2 Diabetes, T2DadjBMI: Type 2 Diabetes adjusted for BMI, TC: Total Cholesterol, TG: Triglycerides, DBP: Diastolic Blood Pressure, MAP: Mean Arterial Pressure, PP: Pulse Pressure, SBP: Systolic Blood Pressure, AF: Atrial Fibrillation, HR: Heart rate, EA: Educational attainment)

## Discussion

In this study, we undertook the largest two-sample MR analyses of thyroid function (TSH and FT4 levels) on CAD and stroke risk. Our results show that higher TSH levels within the normal range are associated with lower the risk of stroke, and that this effect is likely mediated by a lower risk of AF. Furthermore, we find a causal association between HD and CAD.

### Higher normal range TSH levels lower the risk of stroke via a lower risk of atrial fibrillation

Ischemic stroke is the most common type of stroke, responsible for approximately 80% of all stroke incidents(41). Several modifiable cardiovascular risk factors for ischemic stroke have been identified, including hypertension, T2D, dyslipidaemia, obesity and AF(42). Observational studies showed that even variation in thyroid function within the normal range is associated with these factors(43), suggesting that minor variation in thyroid function might be a potential risk factor for stroke.

In 2016, a multi-center study including 43,598 participants investigated the association between variation in normal range thyroid function and stroke risk (9). This study showed that higher TSH levels within the normal range were associated with a decreased risk of stroke, while higher FT4 levels within the normal range were associated with an increased risk of stroke. These effects were independent from traditional cardiovascular risk factors including SBP, TC levels, smoking status and T2D (9).

In our study, we investigated whether causal relationships underlie these epidemiological observations. We demonstrate that higher TSH levels within the reference range are causally associated with a decreased risk of stroke, which is mediated by the decreased risk of AF. This is in line with the results of observational studies which have shown that AF is a major risk factor for stroke, increasing stroke risk up to five-fold (44). Our findings are also in line with recent studies showing that subjects with a genetically predicted higher TSH level have a lower risk of AF (40). While multiple pathophysiological pathways might be theoretically involved in the effect of thyroid function on stroke, we demonstrated that AF has a major role in mediating the effect of TSH within the reference range on the stroke risk.

However, we did not observe a causal effect of FT4 levels on the risk of stroke. This may be because genetic variants used as instruments in our study could explain less variance in FT4 than in TSH levels (i.e. 9.4% and 4.8% of the genetic variance in TSH and FT4 levels, respectively)(18) and therefore our study was better powered to detect the causal effect on the stroke risk for TSH. This suggests that the causal effect of minor variation in FT4 levels on the stroke risk is smaller than detectable in our study.

### MR analysis does not support a causal association between normal range thyroid function and the risk of CAD

Observational studies have shown that even thyroid hormone levels within the normal range are associated with subclinical atherosclerosis, as assessed by coronary artery calcification score and the overall risk of adverse atherosclerotic cardiovascular events, including fatal and nonfatal myocardial infarction, other CAD mortality and stroke (10). However, several large population-based studies investigated the association between variation in normal range thyroid function and the CAD risk only, and did not find an association (8, 45–47). Nevertheless, most of these studies have shown associations between normal range TSH and/or FT4 levels and CAD mortality (8, 45–47). While this may suggest an important relation between thyroid function and CAD, observational studies are prone to residual confounding(48). Our MR study does not provide evidence for a causal association between normal range thyroid function and the risk of CAD. However, we cannot exclude that thyroid function affects the CAD risk with a smaller effect than which was detectable in our study. The effect of minor variation in thyroid function on the CAD risk might be potentially mediated by other factors, such as hypertension and dyslipidaemia, which are nowadays widely recognised and treated. Therefore the potential cause-and-effect relationship between thyroid function and the CAD risk might be limited in subjects on lipid-lowering or/and antihypertensive treatment. This would make the effect of minor variation in TSH and FT4 levels on the CAD risk less pronounced in the general population and consequently more difficult to detect in our study.

### Hashimoto’s Disease and CAD risk

Both overt and subclinical hypothyroidism have been associated with an increased risk of atherosclerosis and adverse cardiovascular events, including CAD, in observational studies (3, 4, 49). In our study, we provide evidence supporting a causal association between HD and CAD, which is mediated by BMI. However, it is not clear whether this effect is attributed solely to hypothyroidism or to autoimmunity in general, since many observational studies suggest an increased risk of atherosclerosis and adverse cardiovascular events in patients with other autoimmune disorders (50). Therefore, future studies are needed in order to clarify the underlying pathophysiological relationship between HD and CAD.

### Strengths and limitations of the study

This is a study comprehensively investigating the causal associations between thyroid (dys)function and adverse cardiovascular outcomes, focusing on stroke and CAD. In order to optimize the power of our study, we used summary data from the largest available GWAS on thyroid function recently published by the ThyroidOmics Consortium, which more than doubled the number of variants associated with TSH and FT4 levels (18). We also used summary statistics from the two largest available GWAS meta-analyses on the tested cardiovascular outcomes, both including >500,000 participants (19, 20). While we found evidence of causal association between TSH levels and stroke, we also investigated potential mediators of this effect and proved that this effect is mediated *via* AF.

Given the relatively large number of variants with unclear physiological function included in the MR analyses, it is possible that some of them may still confer pleiotropic effects. However, using multiple variants associated with TSH and FT4 levels should significantly reduce the impact of individual SNPs associated with the outcome through alternative pathways(14).

Furthermore, a higher number of variants are included as instruments, which explain a higher portion of variance in FT4 and TSH levels (i.e. 9.4% and 4.8% of the genetic variance in TSH and FT4 levels, respectively) (18).

Finally, the IVW method can lead to moderately biased estimates for binary outcomes(51). To address this issue, we applied different MR approaches, which led to consistent results.

## Conclusions

We demonstrate that higher TSH levels within the normal range is associated with a lower the risk of stroke, which is mediated by a decreased risk of AF. These results introduce minor variation in normal range thyroid function as a novel modifiable risk factor for stroke. Additionally, we show that the HD effect on CAD is mediated *via* BMI. This is important as thyroid dysfunction affects 5-10% of the general population, and these results pave the way to consider future adjustment of thyroid function within the normal range in managing patients’ risk of stroke. These findings also have clinical implications as they pave the way to consider future adjustment of thyroid function within the normal range in managing patients’ risk of stroke.

## Acknowledgments

This work was supported by the British Heart Foundation (BHF) grant RG/14/5/30893 to P.D. This work was supported by the Exchange in Endocrinology Expertise (3E) program of the European Union of Medical Specialists (UEMS), Section and Board of Endocrinology. This project has been supported by the Exchange in Endocrinology Expertise (3E) program of the European Union of Medical Specialists (UEMS), Section and Board of Endocrinology (A.K.) This research has been conducted using the UK Biobank Resource under Application Number 9922. The MEGASTROKE project received funding from sources specified at http://www.megastroke.org/acknowledgments.html.

## Disclosures

The authors have no competing interest to declare

